# Reactive blue 2 labels protamine in late-haploid spermatids and spermatozoa and can be used for toxicity evaluation

**DOI:** 10.1101/2023.03.06.531276

**Authors:** Satoshi Yokota, Tomohiko Wakayama, Hidenobu Miyaso, Kousuke Suga, Masakatsu Fujinoki, Satoru Kaneko, Satoshi Kitajima

## Abstract

**Background:** Reactive blue 2 (RB2) dye specifically binds to the nuclei of human spermatozoa under weakly alkaline conditions, thus providing a new method to assess sperm quality. However, this technique has not yet been applied to other mammalian species, such as well-established rodent models, which could enable evaluation of the male reproductive toxicity of drug candidates in non-clinical studies.

**Objectives:** We aimed to evaluate the usefulness of RB2 staining in assessing testicular and epididymal sperm toxicity in mice using a busulfan-induced infertility model.

**Methods:** Male C57BL/6J mice were intraperitoneally administered 40 mg/kg of busulfan. After 28 days, the testes and epididymis were collected and stained with RB2 at pH 10. In vitro evaluations were conducted on uncoated glass slides with RB2 mixed with either protamines extracted from the spermatozoa or intracellular protein components from somatic cells without protamines.

**Results:** Following peanut agglutinin (PNA) lectin histochemistry, RB2-positive cells were observed in elongating and elongated spermatids at all stages except for stages IX–XI of the seminiferous epithelium. After busulfan administration, the proportion of RB2-positive germ cells in the seminiferous tubules decreased significantly, and no RB2-positive spermatozoa were found in the caput epididymis of treated mice. Aggregates were observed in the mixture of RB2 dye (pH 10) with protamines but not in the mixture of intracellular protein components without protamines, and this specificity was lost at neutral pH.

**Discussion and Conclusion:** Our study demonstrates that RB2 specifically stains steps 12–16 spermatids, indicating specific binding to protamine expressed in these spermatids. The RB2 staining technique has potential as a biomarker for male reproductive toxicity, allowing for the rapid visualization of protamination.

## Introduction

The International Council for Harmonisation of Technical Requirements for Pharmaceuticals for Human Use guidelines require histopathological evaluation of the effects of a candidate drug using hematoxylin-eosin (HE) staining in non-clinical animal studies, including general toxicity studies as well as male reproductive toxicity studies.^1^ Specifically, testicular toxicity must be evaluated prior to the first clinical trial in humans. If pharmaceutical candidates are shown to have potential testicular toxicity, non-clinical animal testing has to be withdrawn. Although there are no specific recommendations in regulatory guidelines for general toxicity studies using animals regarding the evaluation of male reproductive toxicity, non-clinical testicular toxicity testing could be preferable to detect similar changes that may occur in clinical trials. However, the results of non-clinical animal testing do not necessarily match the effects in humans.^2, 3^ Furthermore, identification and interpretation of chemically induced changes in testis histology require fundamental knowledge of spermatogenesis; however, there is currently no staining technique available to specifically identify male germ cell development using HE staining in complicated histology. Thus, the development of a novel method to specifically stain spermatogenic cells is needed to support the interpretation of the results of histopathological analysis using HE staining.

During the late haploid phase of spermatogenesis (spermiogenesis), mammalian spermatozoa undergo protamination, in which approximately more than 90% of the histones that package DNA in early spermatids are removed from the DNA and are replaced by transition nuclear proteins and finally by protamines (PRMs).^4^ PRMs possess more domains rich in arginine (Arg), a basic amino acid, than histones and transition nuclear proteins to promote sperm head condensation and DNA stabilization by binding to the phosphoric group in DNA.^5^ The importance of PRMs in sperm chromatin condensation and the influence of protein expression on sperm function have been shown in both mouse and human studies.^6–9^ In most mammals, the sperm chromatin is packaged by a single PRM, whereas primates and most rodents, as well as a subset of other placental mammals, express two PRMs: protamine 1 (PRM1) and protamine 2 (PRM2). Alterations in the content of sperm PRMs induce negative effects on sperm concentration, motility, and sperm head morphology in men.^8^ Haploinsufficiency of PRMs in mice results in sperm morphological abnormalities, DNA damage, and decreases in sperm motility.^10^ In particular, PRM2 deficiency has a negative impact on chromatin packaging and sperm head morphology.^7^ Incorrect condensation of sperm chromatin leads to head abnormalities such as larger heads.^11^ These findings suggest that PRM levels could serve as a biomarker of male infertility.^6^

A previous study showed that reactive blue 2 (RB2) dye specifically bound to the nucleus of spermatozoa under weakly alkaline conditions (pH 10), providing a new method for assessing human sperm quality.^12^ At this weakly basic pH, the three sulfate residues in RB2 may bind to the guanidyl groups in Arg through electrostatic interactions, enabling specific staining of PRMs in human spermatozoa.^12^ Based on the chemical structure of RB2, we hypothesized that RB2 specifically binds to PRMs in male haploid germ cells of other mammals under weakly alkaline conditions. It would be useful to apply RB2 staining in non-clinical studies for evaluating male reproductive toxicity in rodent models; however, this technique has not yet been attempted in mammalian species other than humans.

Therefore, the aim of the present study was to examine whether the germ cell specificity of RB2 found in humans is consistent in experimental mammalian models such as mice, which would enable research on male reproductive toxicity in the context of complicated histology. Specifically, we investigated whether PRMs could be visualized by RB2 using an in vitro experiment to examine the usefulness of this novel staining technique for evaluating mouse testicular and epididymal sperm toxicity using a standard mouse model of infertility established by the administration of busulfan.^13–16^

## Materials and Methods

### Animals

Twelve C57BL/6J male mice (CLEA Japan, Tokyo, Japan) aged 5 weeks were maintained under standard laboratory conditions. The mice were housed in ventilated cage systems (Lab Product Inc., Seaford, DE, USA) at 23 ± 2°C under a 12-h light/dark cycle (lights on from 8:00 to 20:00), with food and water provided *ad libitum*. Body weight (21.46 ± 0.9 g) was measured once a week (three mice per cage). This study followed the guidelines established by the Ethical Committee for Animal Experiments of the National Institutes of Health Sciences (NIHS). The animal facility was approved by the Health Science Center for the Accreditation of Laboratory Animal Care in Japan. All experimental protocols in the study were reviewed and approved by the Committee for Proper Experimental Animal Use and Welfare, a peer-review panel established at the NIHS, under experimental approval no. 816.

### Experimental design

After a 1-week habituation, the 12 mice were randomly divided into two groups: the vehicle control group (n = 6) and the busulfan-treated group (n = 6). The vehicle control group received two intraperitoneal (i.p.) injections of vehicle containing 10% dimethyl sulfoxide (DMSO; 031-24051; CultureSure®DMSO; FUJIFILM Wako Pure Chemical Industries, Ltd., Osaka, Japan) in saline (Otsuka Pharmaceutical Co., Tokyo, Japan) with a 3-h interval between injections. Busulfan (B-2635; Sigma-Aldrich, St. Louis, MO, USA) was dissolved in DMSO at a concentration of 10 mg/mL and then gradually added to a nine-times volume of saline (final concentration: 1 mg/mL). Similar to the vehicle control group, the busulfan-treated group also received two i.p. injections (20 mL/kg per injection) of busulfan at a total dose of 40 mg/kg body weight, as previously described.^13, 17^ After 28 days, the reproductive organs of the male mice were collected under 3.5% isoflurane anesthesia. All efforts were made to minimize animal suffering. Each mouse was weighed, blood was collected from the inferior vena cava, and the testes and epididymis were removed. Dissected tissues were fixed for histopathological analysis as described below.

### RB2 staining of spermatids in the testes and of spermatozoa in the epididymis

For histopathological analysis, the testes and epididymis were removed and immersed in a freshly prepared 4% paraformaldehyde phosphate buffer fixative (161-20141; FUJIFILM Wako Pure Chemical Industries, Ltd.) for 48 h. Following fixation, the testes and epididymis were processed using a graded alcohol series, cleared in xylene, and embedded in paraffin. Sections (4-μm thick) were prepared, deparaffinized, and stained with HE using our routine method for overall morphological evaluation. Two serial testis sections were prepared: one was used for visualization of male germ cells expressing protamine according to RB2 staining, and the other was applied for staging of the mouse seminiferous epithelium cycle using peanut agglutinin (PNA) lectin histochemistry, as previously described.^18, 19^ For the former analysis, the testicular and epididymal cross-sections were stained with 0.1% RB2 (catalog no. 12236-82-7; RB2; R115; Sigma-Aldrich) in 100 mM carbonate-bicarbonate buffer (pH 10) for 30 min, and the excess dye was then washed out with the carbonate-bicarbonate buffer. For the latter, the sections were immersed in 20 mM Tris-HCl buffer (pH 9) and heated at 90°C for 30 min for antigen retrieval. After cooling to room temperature, the sections were washed in phosphate-buffered saline, incubated with Alexa Fluor 488-conjugated lectin PNA antibody (L21409, 1:1000; Molecular Probes, Eugene, OR, USA) for 30 min at room temperature to visualize the acrosomes, and then counterstained with Hoechst 33258 (Sigma-Aldrich). Each stage of spermatogenesis was determined to simplify the identification of the cell population, as previously reported.^18, 19^ Briefly, testis sections from the control mice were observed to determine the staging of the seminiferous epithelium cycle. The 12 stages (I–XII) of spermatogenesis in mice were then determined using the combination of PNA lectin histochemistry for the acrosomes of spermatids and Hoechst staining for chromatin in the nuclei, according to the established criteria of stages based on the position, size, and shape of acrosomes.^18, 19^

The slides were mounted using ProLong Diamond Antifade Mountant (P36961; Thermo Fisher Scientific, Kyoto, Japan) prior to microscopic examination. To quantify the proportion of RB2-positive germ cells in the seminiferous tubules, 200 seminiferous tubule sections were observed in two tissue sections obtained from two discrete portions of a testis from a single mouse. We evaluated the two tissue sections per mouse (n = 6 per group) using a microscope (BX51; Olympus Co., Tokyo, Japan) equipped with cellSens imaging software (Olympus Co., Tokyo, Japan).

### Isolation of spermatozoa from the caput or cauda epididymis and RB2 staining

For isolation of spermatozoa, the caput or cauda epididymis from each animal was cut using a surgical blade and then minced using small scissors in 10 mM HEPES-buffered TYH culture medium (119 mM NaCl, 4.8 mM KCl, 1.7 mM CaCl_2_, 1.2 mM KH_2_PO_4_, 1.2 mM MgSO_4_, 25 mM NaHCO_3_, 5.6 mM glucose, 1.0 mM sodium pyruvate, and 4 mg/mL bovine serum albumin; pH 7.4). A small sample of the sperm suspension from the caput or cauda epididymis was spread onto a glass slide using a centrifugal auto-smear technique (Cyto-Tek; Sakura, Tokyo, Japan). The spermatozoa were then fixed with methanol for 5 min and stained with 0.1% RB2 in 100 mM carbonate-bicarbonate buffer (pH 10) for 30 min. The slides were rinsed with carbonate-bicarbonate buffer to remove any excess dye. They were then mounted using ProLong Diamond Antifade Mountant (P36961; Thermo Fisher Scientific) prior to microscopic examination. We observed 200 spermatozoa per mouse (n = 6 per group) using the BX51 microscope equipped with cellSens imaging software.

### Analysis of motility of spermatozoa

For motility analysis, cauda spermatozoa were collected according to our previously reported method with minor modifications.^19, 20^ Briefly, the cauda epididymis was dissected, and the connective tissue, fat pad, muscles, and vas deferens were removed. For sampling of spermatozoa, the cauda epididymis from each animal was cut using a surgical blade and then minced using small scissors in 10 mM HEPES-buffered TYH culture medium, as described above. The suspensions of spermatozoa were left to disperse and swim up for 15 min on a warming tray at 37°C. The suspension was then filtered through a 40-μm nylon filter (PP-40N; Kyoshin Rikoh, Tokyo, Japan) to remove any undigested tissue fragments, and cauda spermatozoa were collected to evaluate motility. The motile spermatozoa were incubated for 2 h at 37 °C to allow for hyperactivation.

The percentage of motile or hyperactivated spermatozoa was measured on a 37°C heated stage of a constant-temperature unit (MP-10; Kitazato Supply Co., Shizuoka, Japan), according to a previously described method.^20^ Sperm motility was recorded using a phase-contrast microscope (CX43; Olympus, Tokyo, Japan) with Moticam 1080 (Shimadzu Rika Co., Tokyo, Japan). Using a 10× phase-contrast objective, each field was recorded for 30 s, and the numbers of total spermatozoa and motile and/or hyperactivated spermatozoa were manually counted in three random fields for each sample. The analyses were performed in a blinded manner. Motile spermatozoa that exhibited asymmetric and whiplash flagellar movements and a circular and/or octagonal swimming locus were defined as hyperactivated.^20, 21^ The percentage of motile spermatozoa was calculated as the number of forward motile and sub-motile spermatozoa or hyperactivated spermatozoa/total number of spermatozoa × 100 (%).

### *In vitro* experiment of RB2 binding

RB2 dye in 100 mM carbonate-bicarbonate buffer (pH 10) was mixed with human spermatozoa PRM (Briar Patch Biosciences, CA, USA) or mouse synthetic PRM1 or PRM2 (Briar Patch Biosciences) on non-coated glass slides. After the reaction, the glass slides were mounted using a mountant and observed under a microscope. As a negative control, bovine serum albumin (BSA; 23209; Pierce™ BSA standard ampules, 2 mg/mL; ThermoFisher Scientific, Waltham, MA, USA) and an extracted intracellular protein from the human HeLa cell line (from the Japanese Collection of Research Bioresources Cell Bank) were mixed with RB2 dye (pH 10). The HeLa cells were cultured in Dulbecco’s modified Eagle’s medium (D5030, Sigma-Aldrich) supplemented with 10% fetal bovine serum (10099, Thermo Fisher Scientific) and 100 μg/mL kanamycin (420311, Sigma-Aldrich). The cells were lysed with sodium dodecyl sulfate (SDS) lysis buffer (100 mM Tris-HCl at pH 8, 10% glycerol, and 1% SDS) to extract the intracellular proteins.

### Statistical analysis

Data are presented as mean ± standard deviation (GraphPad Prism version 9, GraphPad Software Inc., San Diego, CA, USA). Welch’s *t*-test was used to detect significant differences between the control and busulfan-treated groups (GraphPad Prism version 9). Statistical significance was assessed at P < 0.05.

## Results

### Body and organ weights

There were no significant differences in body weight between the control and busulfan-treated mice (control; 25.3 ± 0.7 g, busulfan-treated; 24.9 ± 0.4 g). Images of the gross appearance of the testes and epididymis in both the control and busulfan-treated mice are shown in **Supplemental Figure 1A–D**. The absolute weights of the testes (left: control = 104.6 ± 3.1 mg, busulfan-treated = 32.6 ± 2.7 mg, P < 0.001; right: control = 108.7 ± 7.8 mg; busulfan-treated = 31.5 ± 3.9 mg, P < 0.001) and the epididymis (left: control = 36.4 ± 3.3 mg, busulfan-treated = 25.3 ± 2.0 mg, P < 0.001; right: control = 36.6 ± 3.2 mg, busulfan-treated = 25.6 ± 2.2 mg, P < 0.001) were significantly lower in the busulfan-treated mice than in the control mice. In particular, the caput epididymis of busulfan-treated mice was significantly smaller than that of the control mice (**Supplemental Figure 1D**).

### RB2 specifically stains male haploid germ cells expressing PRMs

Based on the shape of the acrosomes of the round and elongating spermatids (steps 1–12), the corresponding staging of the mouse seminiferous epithelium cycle was determined (stages I–XII) (**Supplemental Figure 2**). After identification of the germ cell population by staging of the mouse seminiferous epithelium cycle (**Figure 1A, C, E, G, I, K, M, O, Q, S, U, W**), using adjacent testis sections from the control mice, RB2-positive cells were observed in steps 12 to 16 of elongating and elongated spermatids in stages XII and I to VIII, but not in stages IX to XI of the seminiferous epithelium cycle (**Figure 1B, D, F, H, J, L, N, P, R, T, V, X**).

**Figure 1.**
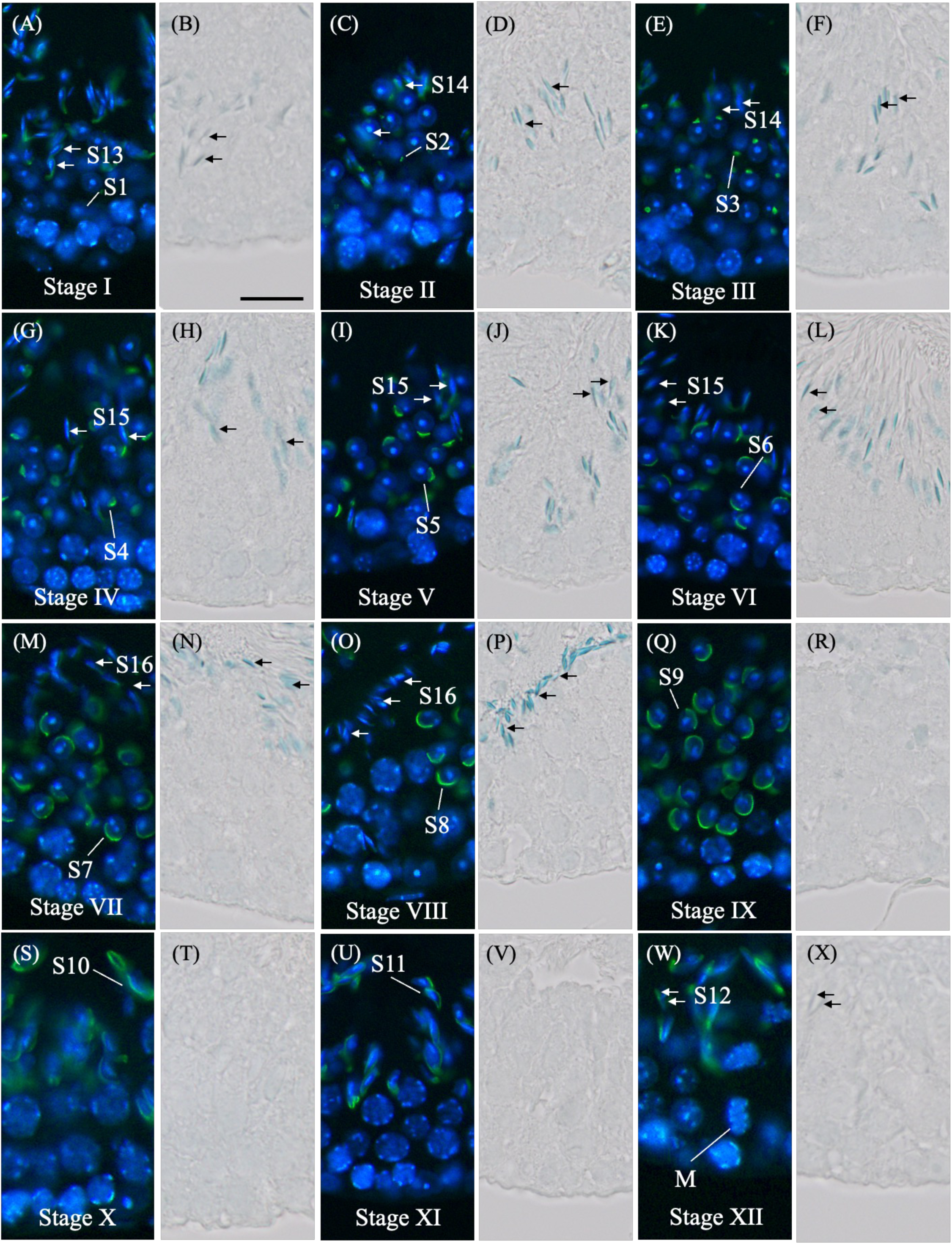
Distribution of reactive blue 2 (RB2)-positive male germ cells in the seminiferous tubules. (A, C, E, G, I, K, M, O, Q, S, U, W) Determination of stages (I–XII) and identification of cell types. The nuclei of mouse control testis sections were stained with Hoechst 33258 (blue) and the acrosomes were stained with PNA lectin histochemistry (green). (B, D, F, H, J, L, N, P, R, T, V, X) Haploid male germ cells were stained with RB2 dye. M, spermatocytes undergoing the first meiotic division; S1–16, steps 1–16 spermatids. White lines: S1–11. White arrows: S12–16. Black arrows: RB2-positive S12–16 spermatids. Scale bars: 20 μm.

Quantitative histological analysis showed that busulfan-treated mice had approximately two times as many seminiferous epithelium tubules as those in the control, with loss of steps 12 to 16 elongating and elongated spermatids (**Figure 2A–G**).

**Figure 2.**
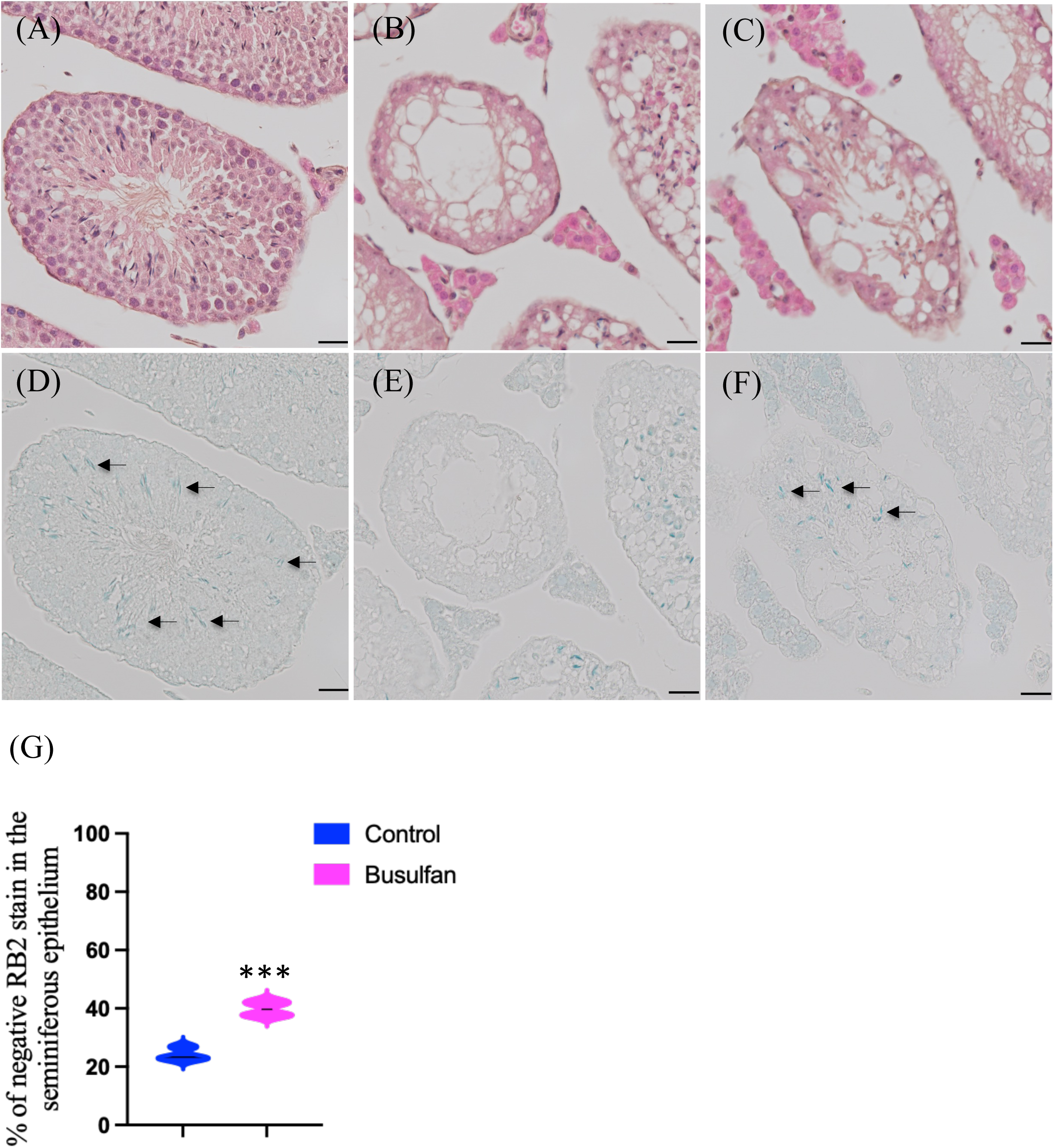
Representative photographs of hematoxylin and eosin (HE)- or reactive blue 2 (RB2)-stained testicular cross-sections from control and busulfan-treated mice. The incidence of RB2-positive spermatids in the seminiferous tubules was examined via light microscopy. (A) HE- or (D) RB2-stained sections from the testes of control mice. (B, C) HE- or (E, F) RB2-stained sections from the testes of busulfan-treated mice. Black arrows: RB2-positive spermatids. (G) Violin plot of the percentage of RB2-positive spermatids in the seminiferous tubules from control (200 tubules/section, n = 6) and busulfan-treated (200 tubules/section, n = 6) mice. Violin plots depicts the frequency distribution of numerical data. Asterisks indicate significant differences between the control and busulfan-treated groups (***P < 0.001). Black arrows: RB2-positive spermatids. Scale bars: 20 μm.

### RB2 staining to evaluate epididymal sperm toxicity

The percentage of motile spermatozoa in busulfan-treated mice was significantly lower than that in control mice (P < 0.01) (**Figure 3A**). The percentage of hyperactivated spermatozoa in busulfan-treated mice also showed a significant decrease compared with that in the control group (P < 0.01) (**Figure 3B**). Swim-up spermatozoa collected from the cauda epididymis of both control (**Figure 3C**) and busulfan-treated (**Figure 3D**) mice were stained with RB2. Spermatozoa collected from the caput epididymis in control mice were also stained with RB2 (**Figure 3E**), whereas those of busulfan-treated mice showed no positive staining (**Figure 3F**).

**Figure 3.**
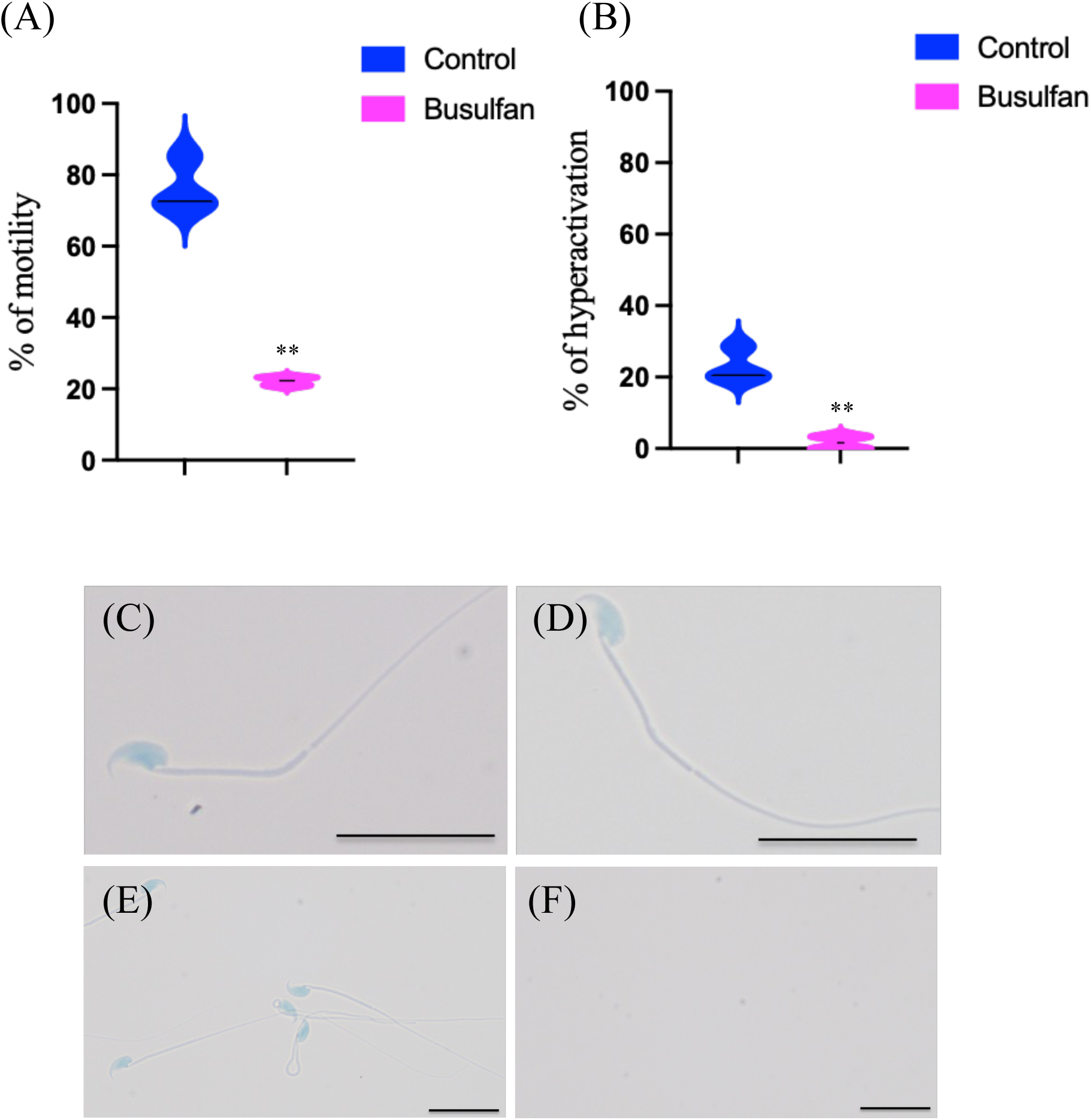
Sperm parameters in the control and busulfan-treated mice. Violin plots showing the percentage of (A) motile or (B) hyperactivated spermatozoa in the control (n = 6) and busulfan-treated (n = 6) mice. Reactive blue 2 (RB2)-stained spermatozoa from the cauda epididymis in the (C) control and (D) busulfan-treated mice. RB2 staining of spermatozoa from the caput epididymis in the (E) control mice and (F) busulfan-treated mice. **P < 0.01 vs. control.

RB2-positive germ cells were detected in the caput epididymis of the control mice (**Figure 4A, C, E, G**), but not in the caput epididymis of busulfan-treated mice (**Figure 4B, D, F**), matching the observations of the spermatozoa collected from the caput epididymis (**Figure 3E**).

**Figure 4.**
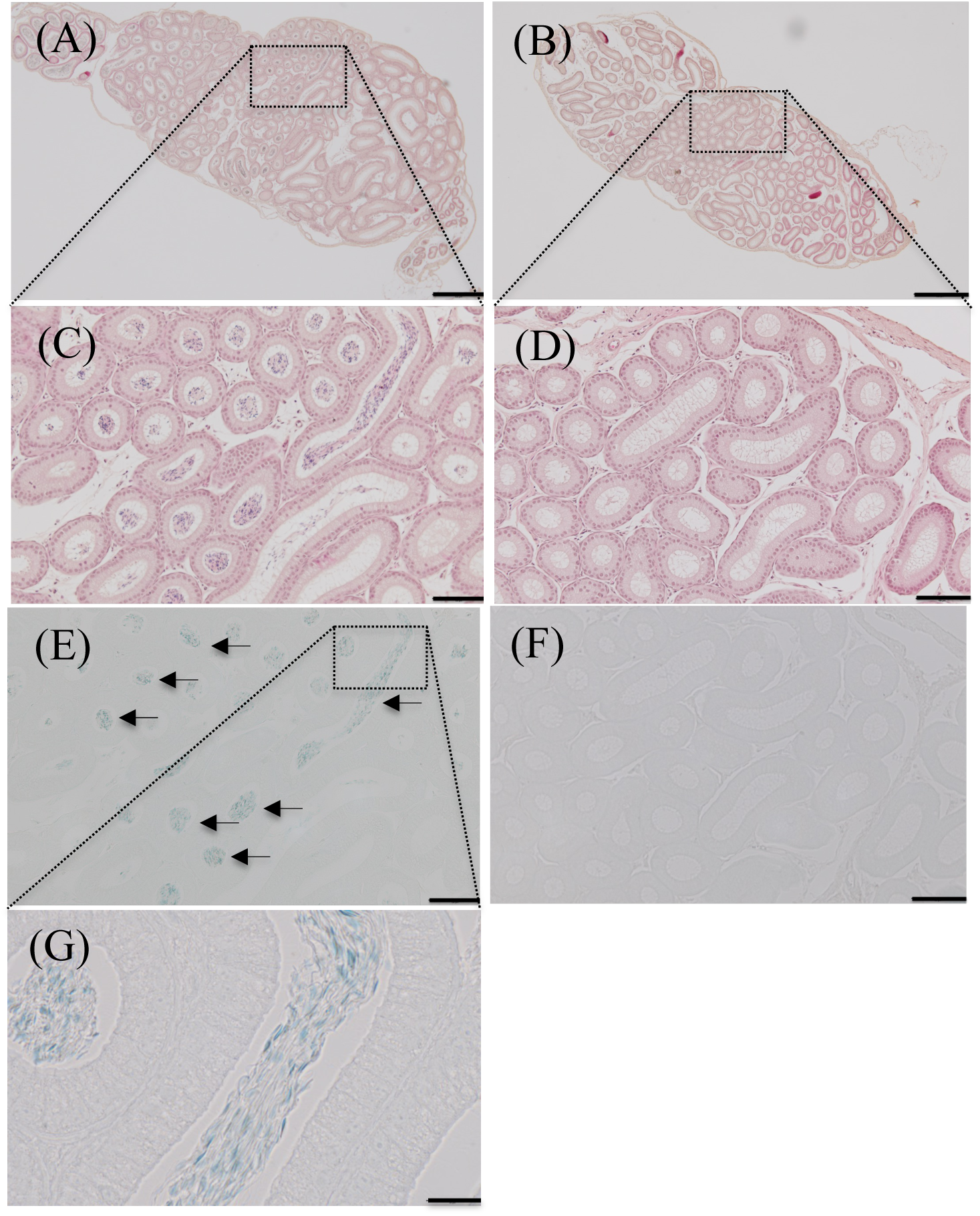
Representative photographs of hematoxylin (HE)- and reactive blue 2 (RB2)-stained caput epididymal cross-sections from control and busulfan-treated mice. Low-magnification images of the morphology in the caput epididymis of (A) control and (B) busulfan-treated mice. High-magnification images of the morphology in the caput epididymis of (C) control and (D) busulfan-treated mice. RB2-positive spermatozoa in the caput epididymis of (E) control mice adjacent to the HE-stained section (C), with no RB2 staining detected in the (F) busulfan-treated group adjacent to the HE-stained section (D). (G) RB2-positive spermatozoa in the caput epididymis of the control under the highest magnification (1000×). Black arrows: RB2-positive spermatozoa. Scale bars: (A, B) 500 μm, (C–F) 100 μm, (G) 20 μm.

### pH-dependence of RB2 specificity for PRMs

Aggregates were observed in the mixture of RB2 dye (pH 10) with both human spermatozoa PRMs and mouse synthetic PRMs (**Figure 5A–F**) but were not detected in the mixture with BSA and the extracted intracellular proteins from HeLa cells (**Figure 5G–I**). In addition, this PRM specificity of RB2 was lost at neutral pH in which histones of the somatic cells were positively stained (**Figure 5J**).

**Figure 5.**
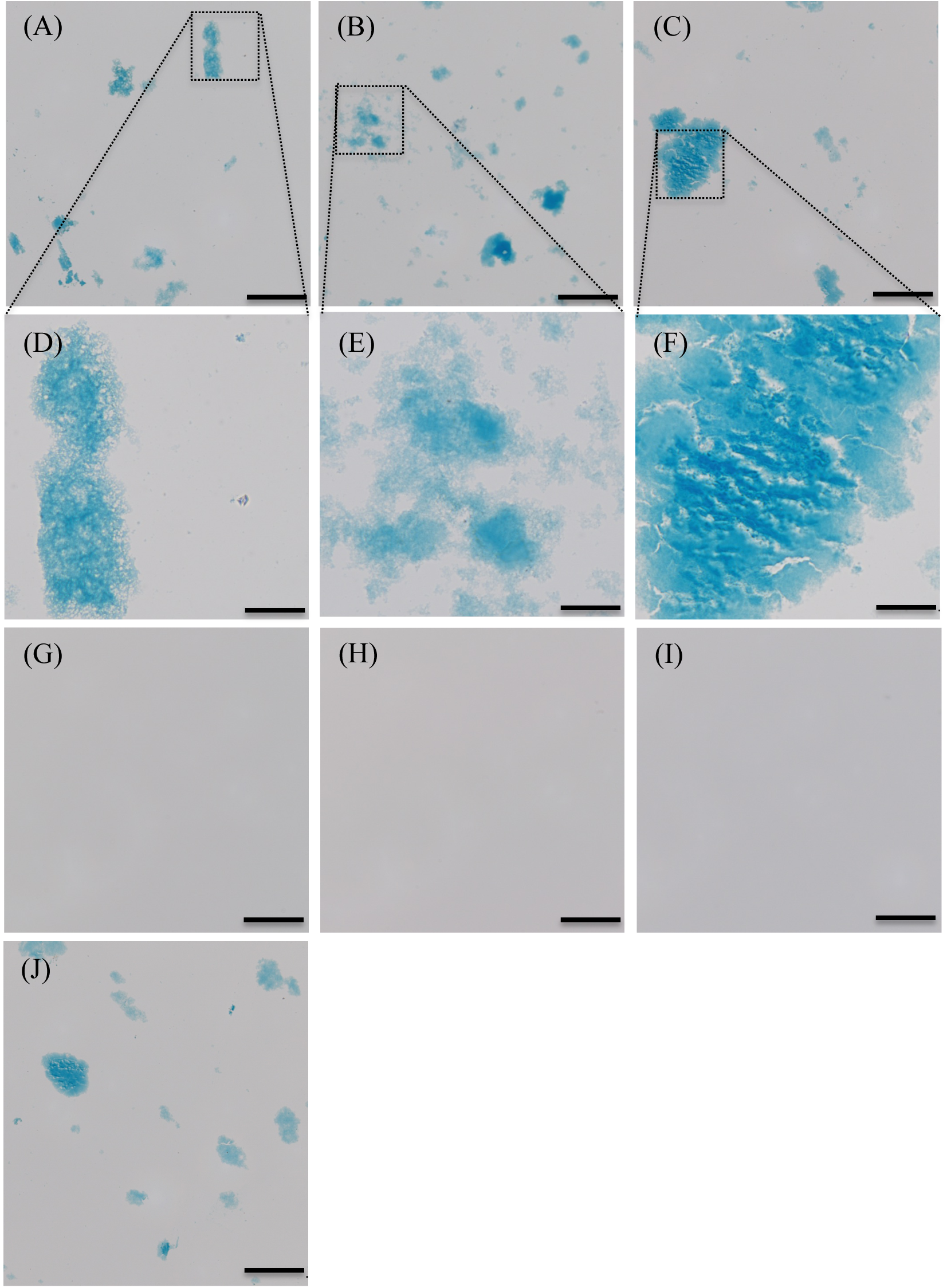
Representative photographs of the mixture of reactive blue 2 (RB2) dye and proteins. Aggregates detected in (A, D) human protamine, (B, E) mouse synthetic protamine 1, and (C, F) mouse synthetic protamine 2 standards. (G, H, I) No aggregates were detected in the RB2 dye only (G), bovine serum albumin standard (H), and intracellular proteins (without protamines) extracted from HeLa cells (I). (J) However, specificity of RB2 was lost at neutral pH in the mixture of RB2 dye and intracellular proteins (without protamines) extracted from HeLa cells. Scale bars: (A–C, G–I) 100 μm, (D–F) 20 μm.

## Discussion

Histopathology of the testes is the best non-clinical endpoint for the evaluation of testicular toxicity. Expertise in spermatogenesis is necessary to identify and interpret chemically induced side effects on testis histology. If a drug candidate compound affects male germ cell development, it should not be further evaluated for drug development. However, the results of non-clinical animal testing may not necessarily align with the effects found in humans.^2, 3^ An ideal evaluation system for non-clinical toxicological studies of drug development would accurately and easily determine testicular toxicity. However, there is currently no available staining technique to identify male germ cell development, specifically in cases of complicated histology.

Kaneko et al.^12^ demonstrated that RB2 dye at pH 10 specifically stained the nucleus of human spermatozoa without staining other cellular components of the human seminal fluid, providing a new method to evaluate sperm quality. The present study further demonstrates that this RB2 staining technique is also applicable to other mammalian species, such as well-established rodent models used for evaluation of the testicular toxicity of drug candidates in pre-clinical studies, providing a novel staining technique for visualizing the nuclei of male mouse haploid germ cells. This demonstration that both human and rodent spermatogenic cells can be visualized by RB2 staining may be useful for extrapolation of the effects of pharmaceuticals on the testes and epididymis of rodents to humans in a non-clinical toxicity test under conditions of limited time and resources.

In mice, the 12 stages (stages I–XII) of spermatogenesis were determined using the combination of PNA lectin histochemistry for the acrosomes of spermatids and Hoechst staining for the chromatin in the nuclei, according to the established criteria of stages based on the position, size, and shape of acrosomes, which is a common method to clearly identify steps 1–16 spermatids.^18, 19^ With this approach, we found that RB2 staining could only visualize steps 12–16 haploid elongating and elongated spermatids, but not spermatogonia, spermatocyte, and steps 1–11 spermatids at pH 10; these findings match those for human spermatozoa.^12^ Primates and most rodents, as well as a subset of other placental mammals, express two PRMs: PRM1 and PRM2.^6^ Both genes are transcribed in round spermatids (steps 7–9 in mice)^22, 23^, and the mRNA is then stored in the form of cytoplasmic ribonucleoprotein particles until their translation in elongating and elongated spermatids (steps 12–16).^24, 25^ Following transcript storage, mouse *Prm1* mRNA seems to be translated earlier (step 12) than *Prm2* mRNA (steps 14, 15).^26^ Therefore, our results suggest that RB2 stain specifically binds to PRMs expressed in the steps 12–16 haploid spermatids in mouse seminiferous tubules.^24–26^

Aggregates were detected in the mixture of RB2 dye (pH 10) with both human sperm PRM- and mouse synthetic PRM reference standards, but not with BSA standard or intracellular proteins without PRMs extracted from the lysis of somatic cells (HeLa cells). These results suggest that a certain molecular component of PRMs can chemically interact with a component of RB2. Coomassie brilliant blue (CBB) staining is a widely used method for the routine visualization of various proteins such as BSA, which binds via ionic bonds and van der Waals forces. However, CBB staining is not specific for a certain protein, requiring an additional protein separation step for identification, such as polyacrylamide gel electrophoresis.^27^ The present study demonstrates the effectiveness of RB2 dye as a novel, rapid, and specific staining technique for visualizing PRMs without the need for protein separation, which could have significant implications for reproductive toxicology and andrology.

We further explored the mechanism by which RB2 dye stains PRMs. We hypothesized that the unique binding specificity of RB2 with PRMs indicates an electrostatic ionic interaction. Lysine (Lys) and Arg are two positively charged amino acids in proteins that have high aqueous dissociation constants (∼10.5 for lysine^28^ and ∼12.5 for Arg^29^), indicating a strong propensity to carry a positive charge at neutral pH. In contrast, at pH > 10, Arg is a positively charged amino acids, whereas Lys is a neutral residue.^28, 29^ Therefore, the negative charge of the three sulfate residues in RB2 could ionically combine with the guanidyl group in Arg at pH 10, but not with the neutral chemical components of lysine. The present study demonstrated that RB2 specifically stained step 12–16 spermatids, in which the majority of PRMs are translated, at pH 10, as previously reported.^24–26^, whereas the specificity of RB2 staining was lost at neutral pH when both Lys and Arg are positively charged. This pH-dependent specificity could be due to differences in the Arg content among the basic proteins, histones, transition proteins, and PRMs. PRMs are typically short basic proteins with a high number of positively charged Arg-rich residues, allowing the formation of a highly condensed complex with the paternal genomic DNA, which has a strong negative charge.^30–33^ The higher Arg content thus allows PRMs to form more stable complexes with DNA than other basic proteins such as histones and transition proteins.^4^ Lys and Arg account for approximately 20–30% of the amino acids in histones, whereas Arg accounts for 70% of the amino acids in PRMs.^4^ Thus, the higher content of Arg in the PRMs of the step 12–16 spermatids could explain the specificity of the novel RB2 staining technique under weakly alkaline conditions (pH 10).

Finally, with respect to the technical application of this method for male reproductive toxicity evaluation, we demonstrated that the percentage of RB2-positive spermatids in the seminiferous tubules of mice was decreased following busulfan administration. In addition, no RB2-positive spermatozoa were detected in the caput epididymis of busulfan-treated mice, whereas spermatozoa of the cauda epididymis were stained with RB2 in both control and busulfan-treated mice. These results suggest that busulfan administration during spermiogenesis and sperm maturation decreases the abundance of PRMs to cause male infertility. Indeed, busulfan treatment significantly decreased the percentage of motile or hyperactivated spermatozoa, as a common index of male fertility. These results suggest that the RB2 staining technique can offer a rapid assessment of the male reproductive toxicity of a drug, such as spermatogenic dysfunction and/or impairment of sperm maturation, with simple visualization of the protamination of male haploid germ cells, providing useful and supportive data to complement the results of conventional HE staining.

The proposed method offers a more convenient and rapid tool for the visualization of PRMs compared with immunohistochemical analysis using anti-PRM antibodies, which will be useful not only for the field of reproductive toxicology but also in general research in the fields of reproductive medicine and andrology to improve the diagnosis of infertility in both humans and rodents. Altered levels of PRMs may result in increased susceptibility to injury of the spermatozoa DNA, causing infertility or poor outcomes in assisted reproduction.^34^ In general, mammalian reproduction relies on the successful transport of the paternal genome to the oocyte via motile spermatozoa, and the formation of functional spermatozoa requires a specific and well-orchestrated differentiation process in the testes.^35^ PRMs have been suggested to be important for remodeling of the chromatin complex following the elongation and condensation of the spermatogenic cell nucleus into a hydrodynamic shape.^34, 36^ Paternal chromatin is completely remodeled from nucleo-histones (nucleosomes) to nucleo-protamines during spermiogenesis to prepare for fertilization.^6, 34^ In clinical studies, PRM deficiency was found to be related to the exacerbation of sperm quality and to cause DNA fragmentation.^37^ Thus, application of spermatozoa with abnormalities in PRM abundance during assisted reproductive technology raises serious concerns for the developing embryo.^38^ Therefore, altered PRM abundance in spermatozoa could serve as an important biomarker of male infertility. In this regard, the use of RB2 as a novel and specific staining technique for PRMs could enable the rapid evaluation of protamination during spermiogenesis as supportive data for histopathological analysis.

Another major finding of the present study was that RB2 staining did not detect any spermatozoa in the caput epididymis within 28 days after the administration of busulfan to mice at 40 mg/kg body weight. Spermatogenesis is a complex process in which many spermatozoa are produced from a relatively small pool of spermatogonial stem cells, which occurs during the entire reproductive life of males.^35^ To the best of our knowledge, most studies on busulfan have largely focused on their testicular toxicity. As an alkylating agent, busulfan causes DNA damage by cross-linking DNA, and was reported to specifically kill A1 spermatogonia (a type of spermatogonial stem cell) as a side effect.^39^ In mice, busulfan administered at 40 mg/kg body weight caused complete depletion of A1 spermatogonia up to the pachytene spermatocytes in stage VIII within 12 days.^40^ Since the proliferation and differentiation of A1 spermatogonia into elongated spermatids takes approximately 34.4 days in the testes^35^, the spermatozoa in the caput epididymis were not considered to be triggered by damage to the A1 spermatogonia within 28 days after busulfan administration. In general, epididymal sperm toxicity evaluation may be more challenging than the evaluation of testicular toxicity. The epididymis is a single, highly convoluted duct lined by a complex pseudostratified epithelium consisting of multiple cell types, including principal cells, basal cells, clear cells, apical cells, and halo cells.^41^ The number and appearance of individual cell types also vary between the segments of the epididymis (i.e., initial segment, caput, corpus, and cauda). The present study demonstrated that RB2 is a specific dye for staining the spermatozoa and could therefore increase the throughput of epididymal toxicity evaluation.

In conclusion, we have demonstrated a novel methodology for visualizing male haploid germ cells using RB2 dye (see **Figure 6**), which provides an important basis for evaluating the effects of pharmaceuticals such as anticancer drugs on testicular and epididymal toxicity. However, this study had some limitations. The data obtained were insufficient for time-course evaluation. Spermatogenesis is a continuous, cyclical, and synchronized process that occurs in the epithelium of the seminiferous tubules of the testes, spanning approximately 34.4 days in mice and 74 days in humans.^35^ In addition, sperm maturation requires approximately 10 days in both mice and humans.^35^ Chemicals may affect the testes and epididymis at any time point throughout these cycles. Therefore, it is difficult to predict precisely when and where testicular and epididymal toxicity may occur, which requires investigation at multiple time points to further assess the potential recovery from drug toxicity, which is particularly important for anticancer drugs. Thus, further studies are needed to evaluate the time-course changes in PRMs abundance in histological testes specimens and in the spermatozoa utilizing the RB2 staining method.

**Figure 6.**
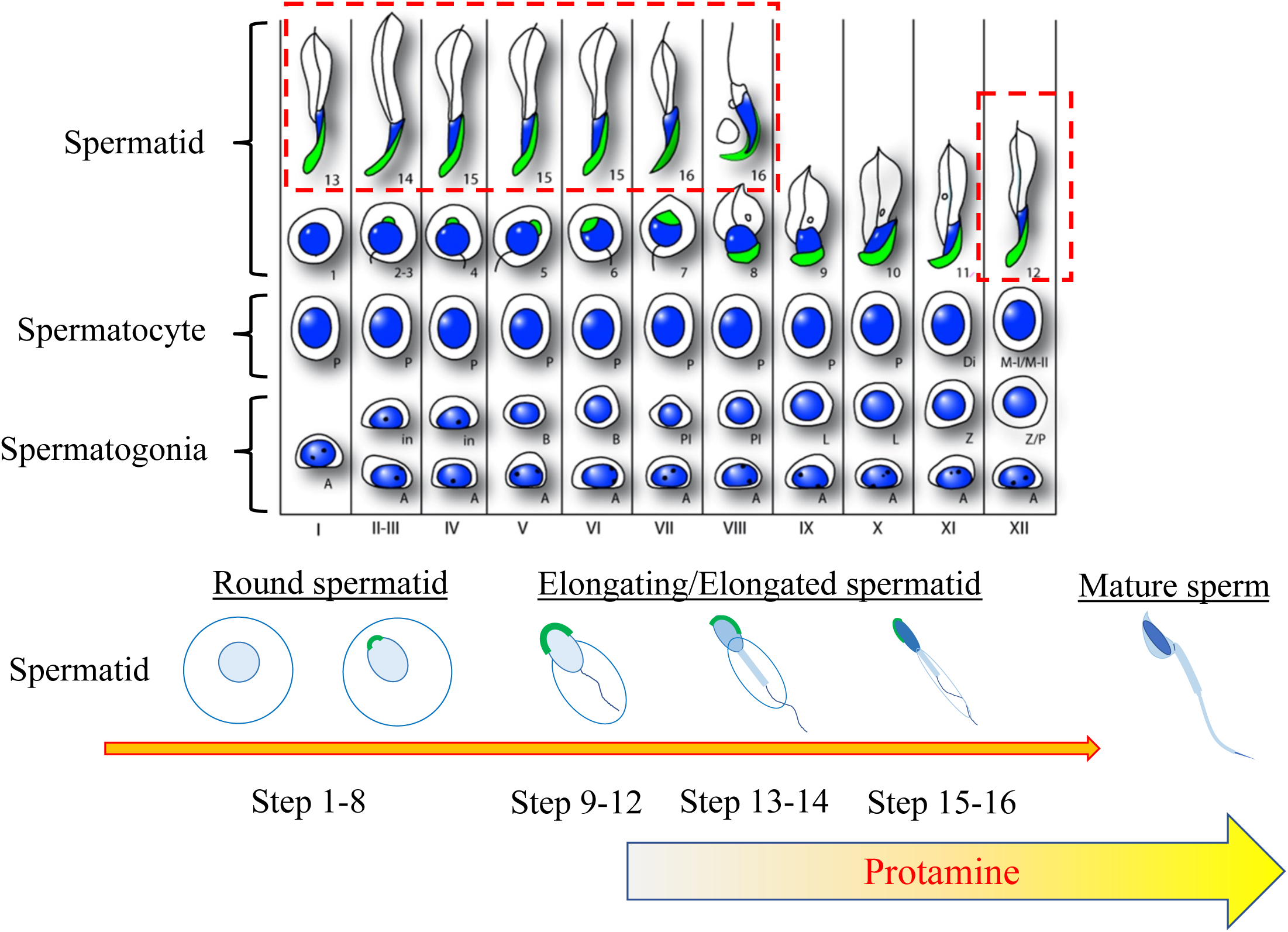
Schematic illustration of reactive blue 2 (RB2)-positive haploid spermatids in the seminiferous tubules.

## Acknowledgments

The authors thank Editage (https://www.editage.jp) for their English language editing service. This work was supported by the Japan Agency for Medical Research and Development (grant number 22mk0101210j0002 to Satoshi Yokota). This manuscript was submitted as a pre-print in the link “https://www.biorxiv.org/content/10.1101/2023.03.06.531276v1”.^42^

## Author contributions

SY conceived and designed the experiments. S Kitajima supervised this study. SY, TW, HM, KS, and MF performed the experiments. SY, TW, S Kaneko, and S Kitajima discussed the results of the data. SY drafted the manuscript. All authors critically revised the article for intellectual content.

## Supplementary description

**Supplemental Figure 1.** Gross appearance of the testes and epididymis in control and busulfan-treated mice.

**Supplemental Figure 2.** Determination of stages I–XII in the mouse seminiferous tubules. The nuclei of mouse control testis sections were stained with Hoechst 33258 (blue) and the acrosomes were stained with PNA lectin (green). (A–C) Stage I, (D–F) Stage II, (G–I) Stage III, (J–L) Stage IV, (M–O) Stage V, (P–R) Stage VI, (S–U) Stage VII, (V–X) Stage VIII, (Y–AA) Stage IX, (AB–AD) Stage X, (AE–AG) Stage XI, (AH–AI) Stage XII. Scale bars: 20 μm.

